# Atypical chemokine receptor 3 (ACKR3) induces the perturbation of rRNA biogenesis: a novel mechanism of colorectal tumorigenesis

**DOI:** 10.1101/2021.09.01.458501

**Authors:** Juan Yang, Ya-Nan Li, Ting Pan, Rong-Rong Miao, Yue-Ying Zhang, Shu-Hua Wu, Xian-Jun Qu, Shu-Xiang Cui

**Author notes:** **Correspondence.** Shu-Xiang Cui. E-mail addresses.

## Abstract

Atypical chemokine receptor 3 (ACKR3), previously known as C-X-C chemokine receptor type 7 (CXCR7), has emerged as a key player in several biologic processes. Its atypical “intercepting receptor” signaling properties have established ACKR3 as the main regulator in pathophysiological processes in many diseases. However, much less is known the underlying mechanisms of ACKR3 in promoting tumorigenesis. We found, in both human and animal model, that activation of ACKR3 promotes colorectal tumorigenesis through the NOLC1-induced perturbations of rRNA biogenesis. As compared with adjacent non-neoplastic tissue, human colonic cancer tissues demonstrated higher expression of ACKR3, and high ACKR3 expression was associated with increased severity of colonic cancer. Villin-ACKR3 transgenic mice demonstrated the characteristics of ACKR3-induced colorectal cancer, showing the nuclear β-arrestin-1-activated perturbation of rRNA biogenesis. Activation of ACKR3 induced nuclear translocation of β-arrestin-1 (β-arr1), leading to the interaction of β-arr1 with nucleolar and coiled-body phosphoprotein 1 (NOLC1). As the highly phosphorylated protein in the nucleolus, NOLC1 further interacted with Fibrillarin, a highly conserved nucleolar methyltransferase responsible for ribosomal RNA methylation, leading to the increase of methylation in Histone H2A, resulting in the promotion of rRNA transcription of ribosome biogenesis. Conclusion: ACKR3 promotes colorectal tumorigenesis through the perturbation of rRNA biogenesis by nuclear β-arr1-induced interaction of NOLC1 with Fibrillarin.

**HIGH LIGHTS:**

- ACKR3 is an atypical G protein-coupled receptor (GPCR)
- ACKR3 promotes colorectal tumorigenesis
- ACKR3 induces nuclear translocation of β-arr1
- Nuclear β-arr1 interacts with NOLC1 to activate Fibrillarin
- Interaction of NOLC1 to Fibrillarin leads to perturbation of rRNA biogenesis

## Introduction

Chemokine receptor ACKR3, formerly named as CXCR7/RDC1, is an atypical G protein-coupled receptor (GPCR) (Bachelerie et al., 2014; Balabanian et al., 2005; Burns et al., 2006). Unlike classical GPCRs, ACKR3 usually fails to activate canonical Gi protein-mediated signaling due to the lack of a conserved DRYLAIV structure (Cancellieri et al., 2013). ACKR3 is once regarded as a scavenger receptor and decoy receptor (Luker et al., 2012; Meyrath et al., 2020). During the pathophysiological processes, ACKR3 could be activated by endogenous ligands CXCL11/I-TAC and CXCL12/SDF-1, resulting in the development of numerous diseases, such as cancers, chronic inflammation, and cardiovascular disorders (Sanchez-Martin et al., 2013). ACKR3 preferentially triggers the β-arrestin-dependent signaling, leading to the internalization *via* the AKT or ERK1/2 signaling pathways (Becker et al., 2019; Li et al., 2019; Rajagopal et al., 2010). At present, still little is known about mechanism of the ACKR3-induced β-arrestin-dependent signaling in tumorigenesis. ACKR3 was found overexpression in many cancers. Activation of ACKR3 has been considered to promote cancer growth through the key processes, such as proliferation, anti-apoptosis, angiogenesis, etc (Miao et al., 2007; Sun et al., 2010; Wang et al., 2008). ACKR3 has thus considered as a potential therapeutic target for the clinical management of cancers. However, these ongoing efforts have been less encouraging in animal models because ACKR3 could participate in a complex signaling network, interacting with additional targets and signaling pathways through various crosstalk and compensatory signaling mechanisms. Further investigating the mechanisms of ACKR3 could help us to fully understand the role of ACKR3 in colorectal tumorigenesis (Salanga et al., 2009).

As a “crosslinker” receptor, ACKR3 is likely to bind with CXCR4, a classic chemokine receptor for CXCL12, to form the ACKR3/CXCR4 heterodimer. In the ACKR3/CXCR4 heterodimer, ACKR3 might function as modulating the CXCR4 signaling to an active and independent signaling receptor (Koch and Engele, 2020). Previously, we revealed a mechanism of ACKR3/CXCR4 heterodimer in promoting colorectal tumorigenesis. ACKR3 was found to modulate or just assist the CXCR4 signaling through inducing β-arr1 recruitment to the nucleus, leading to the increase of histone demethylase JMJD2A (Song et al., 2019). Villin-ACKR3 mice developed more exacerbated colorectal cancer than Villin-CXCR4 mice under the same AOM/DSS condition (Song *et al.*, 2019). Accordingly, ACKR3 might function in tumorigenesis independent of CXCR4 signaling. As yet the mechanism of ACKR3, only known an atypical G protein-coupled receptor, has not been fully investigated.

In the present study, we uncovered a new mechanism of ACKR3 in the promotion of colorectal tumorigenesis through the NOLC1-induced perturbations of rRNA biogenesis. Since β-arr2 has a strong nuclear export signal (NES) in its C terminus, this specific NES excludes it from sustained presence in the nucleus (Meier, 1996; Scott et al., 2002). Thus, in the present study, we identified β-arr1 as a downstream signaling of activated ACKR3 in the pathophysiological processes of colorectal tumorigenesis. The ACKR3-induced nuclear β-arr1 interacted with NOLC1 to form the β-arr1-NOLC1 complex. NOLC1 is a highly phosphorylated nucleolus coiled protein and a binding protein for RNA polymerase I. NOLC1 is essential for small nucleolar riboprotein synthesis and a critical transcription factor for an array of gene transcriptions (Kim et al., 2003; Meier, 1996). In our model, the phosphorylated NOLC1 was further interacted with Fibrillarin, the rRNA methyl-transferase, leading to the promotion of nucleolus Fibrillarin, resulting in the increase of rRNA transcription of ribosome biogenesis through upregulating the methylation in Histone H2A. We envision that furture approaches to treat colorectal cancer should use ACKR3 inhibitors for preventing the ACKR3-activated NOLC1 and Fibrillarin in the nucleolus.

## Results

### High expression of ACKR3 in human CRC specimens

To investigate the role of ACKR3 in colorectal cancer (CRC), we firstly analyzed the data of ACKR3 gene issued in the Oncomine database. An analysis of a Hong Colorectal dataset indicated that the mRNA levels of ACKR3 were significantly higher in colorectal cancers than in normal tissues (P = 7.50E-5, Fold Change = 2.427) (Fig 1A). Importantly, the increased ACKR3 was associated with the progression of clinical stages of colorectal cancer as indicated in the GEPIA database (Fig 1B). F value = 2.23, Pr (>F) = 0.084. We then performed the immunohistochemistry assay (IHC) to determine ACKR3 levels in human CRC tissues and their paracancerous tissues (Fig 1C). Of the 60 CRC specimens, 57 cases were identified higher ACKR3 expression than their paracancerous tissues (Supplementary, Table S4). Western blotting assay further identified higher levels of ACKR3 in human fresh colonic cancer tissues than their paracancerous tissues (Fig 1D). In vitro cultured cells, the cancer cell lines but not human normal colonic cell line demonstrated higher levels of ACKR3 (Fig 1E).

**Figure 1.**
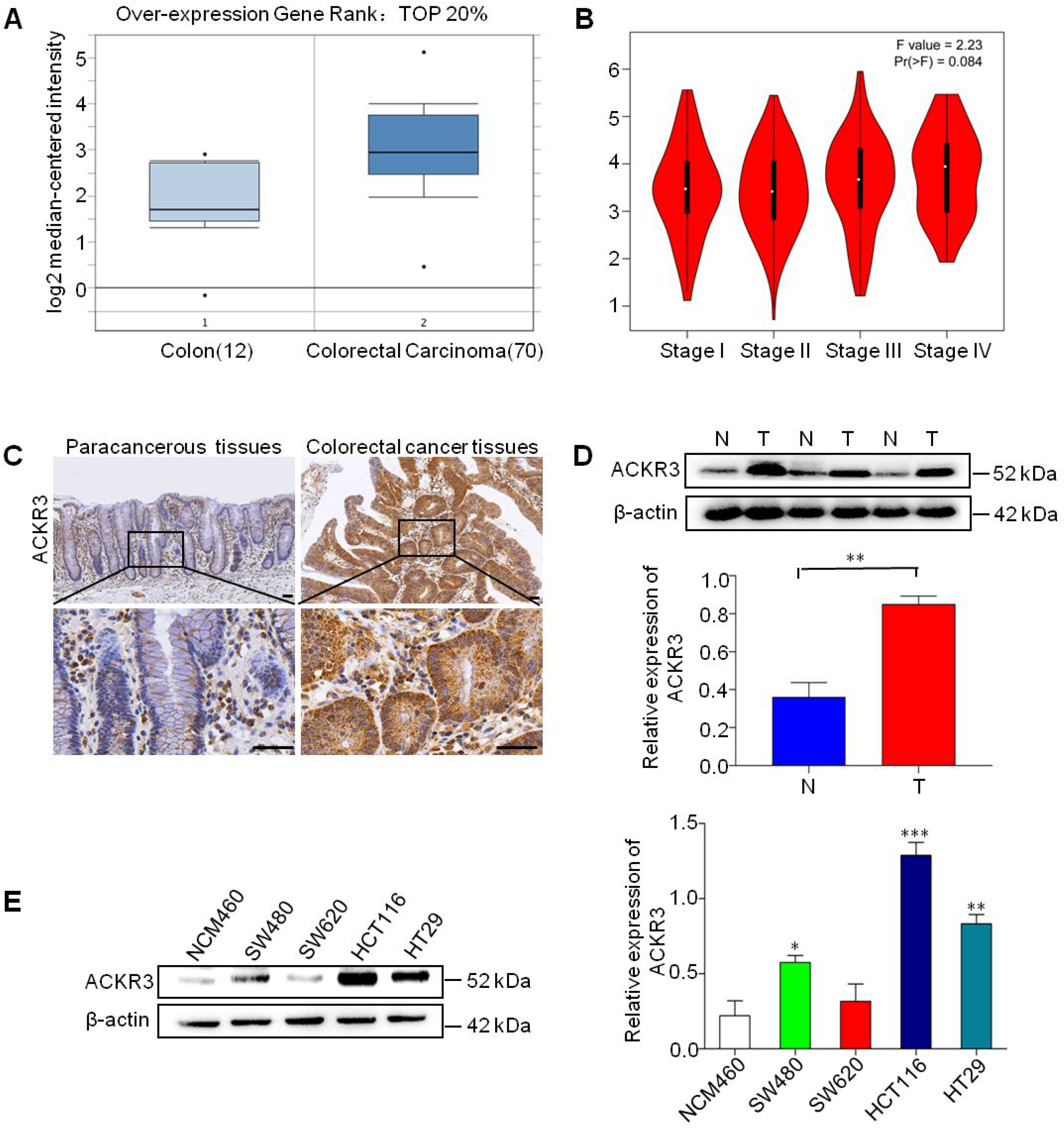
Human CRC specimens were determined high ACKR3 levels. (A) Level of ACKR3 mRNA copy number in colorectal carcinoma and normal tissue in Hong Colorectal dataset. (B) The expression levels of ACKR3 is varied in different clinical stages of colorectal cancers. (C) The comparison of ACKR3 levels in human CRC tissues and paired adjacent non-cancer tissues. Upper images 10x, lower images 40x. The boxed region in each top panel is magnified and shown in the corresponding bottom panel. (10x) Scale bar: 200 μm, (40x) Scale bar: 50 μm. *n* = 20. (D) Western blotting analysis of ACKR3 in the representative (*n* = 20) human CRC tissues (marked by T) and adjacent non-cancer tissues (marked by N). (E) The expression levels of ACKR3 in human colonic cancer cell lines and normal colonic epithelial cell line. Protein level was determined by densitometry and normalized to β-actin (right). \**P* < 0.05, \*\**P* < 0.01, \*\*\**P* < 0.001.

### Increased ACKR3 in intestinal epithelium cells conferred colorectal tumorigenesis in Villin-ACKR3 mice

Villin-ACKR3 mice demonstrated the increased severity of colorectal tumorigenesis as compared to their WT littermates under AOM/DSS induction, showing a significant weight loss (Fig 2A) and an high disease activity index (DAI index) (Fig 2B). AOM/DSS induced shorter colorectal length in Villin-ACKR3 mice showed than WT mice (Fig 2C, D). Importantly, Villin-ACKR3 mice developed more colorectal tumors than WT mice (Fig 2E). These colorectal tumors in Villin-ACKR3 mice demonstrated higher levels of ACKR3 (Fig 2F), with significantly increased CXCL12 (Fig 2G) than those in their WT littermates. Microscopic analysis of colonic tissues showed the exacerbated colonic damage with higher dysplasia in Villin-ACKR3 mice than WT mice (Fig 2H). These symptoms of colorectal tumorigenesis in Villin-ACKR3 mice might associate with the upregulation of signaling of pluripotent potential and the ERK signaling in the intestinal epithelium cells (Supplementary Figure S1 and Figure S2).

**Figure 2.**
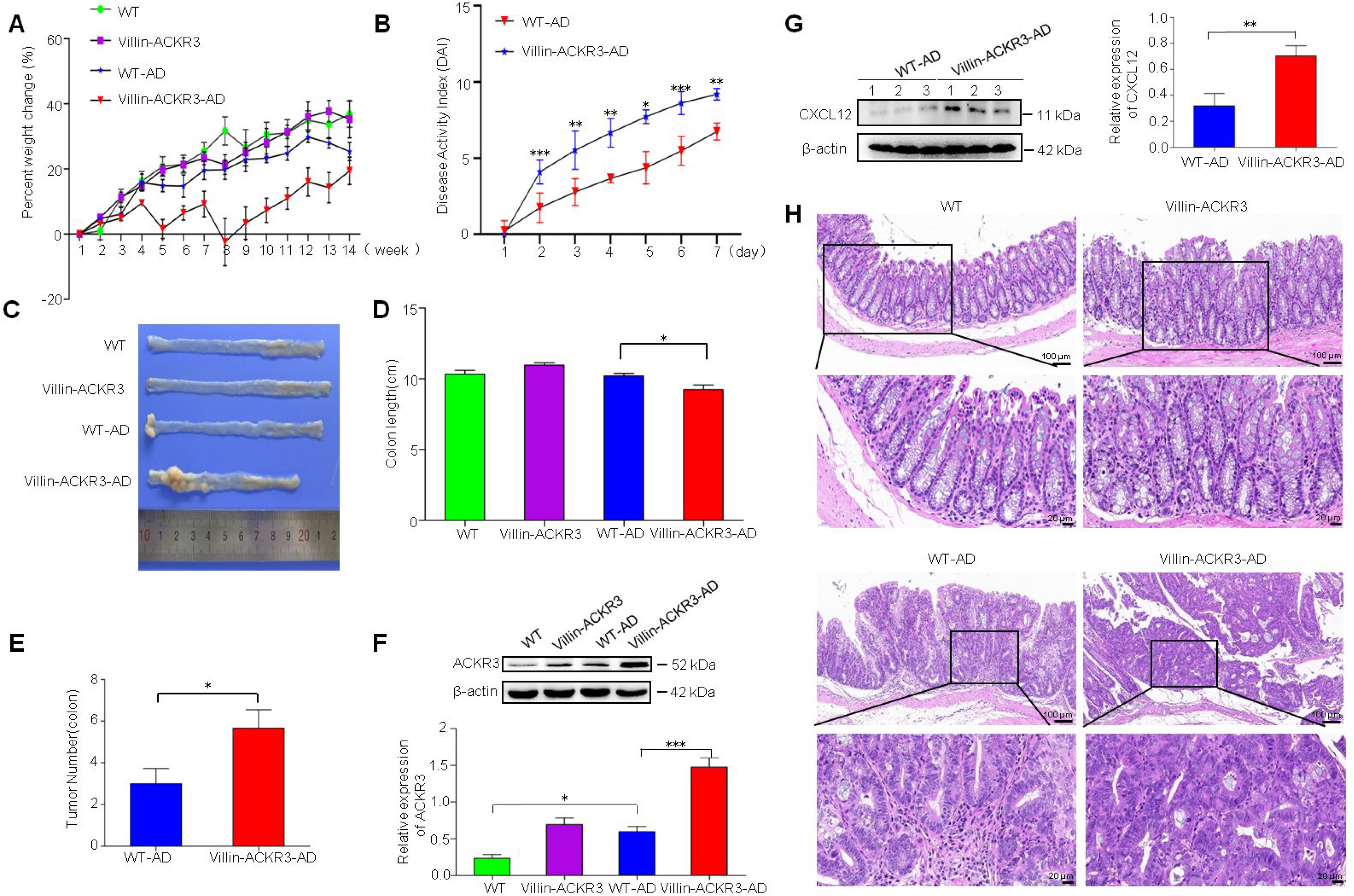
Increased severity of colorectal cancer in Villin-ACKR3 mice. (A) Body weight changes in mice. *n* = 8. (B). Disease activity index score (DAI) during third DSS cycle: WT-AD (Red), Villin-ACKR3-AD (Blue). *n* = 6. (C) Images of colorectal cancer and (D) colorectal length in mice. *n* = 6. (E) Colorectal tumor number in WT-AD, Villin-ACKR3-AD. *n* = 6. (F) Expression levels of ACKR3 in colorectal cancer tissues of WT, Villin-ACKR3, WT-AD, Villin-ACKR3-AD mice; *n* = 3. (G) Western blotting analysis of CXCL12 in CRC tissues of WT-AD and Villin-ACKR3-AD mice. *n* = 3. 1, 2, 3 represented the samples from three mice. (H) H&E-stained colorectal cancer tissues in mice. Upper images 10x, lower images 40x. The boxed region in each top panel is magnified and shown in the corresponding bottom panel. (10x) Scale bar: 100 μm, (40x) Scale bar: 20 μm. *n* = 3. \**P* < 0.05, \*\**P* < 0.01, \*\*\**P* < 0.001. WT-AD: wild mice exposed to AOM/DSS. Villin-ACKR3-AD: Villin-ACKR3 mice exposed to AOM/DSS.

### ACKR3 induced β-arr1 translocation into the nucleus to interact with NOLC1

As the down-stream signal of GPCR, β-arr1 might respond to the activated ACKR3. Human colon cancer cells HCT116 exposed to CXCL12 to activation of ACKR3 demonstrated an increase of β-arr1 in the nucleus with time dependent manner (*P* < 0.05 *vs.* 0 h) (Fig 3A). Since CXCL12 is the ligand of both ACKR3 and CXCR4, we thus used AMD3100, the specific CXCR4 inhibitor, to prevent action of the CXCL12-activated CXCR4 on ACKR3, and then analyzed the expression of β-arr1 in the nucleus. An increase of β-arr1 had remained in the nucleus in the absence of CXCR4 (Fig 3B). Nuclear β-arr1 was clearly seen in the immunofluorescence staining cells (Figure 3C). Nuclear β-arr1 was also seen in colorectal cancer of Villin-ACKR3 mice (Fig 3D). Reversely, silencing of ACKR3 significantly prevented the translocation of β-arr1 into the nucleus (Fig 3E).

**Figure 3.**
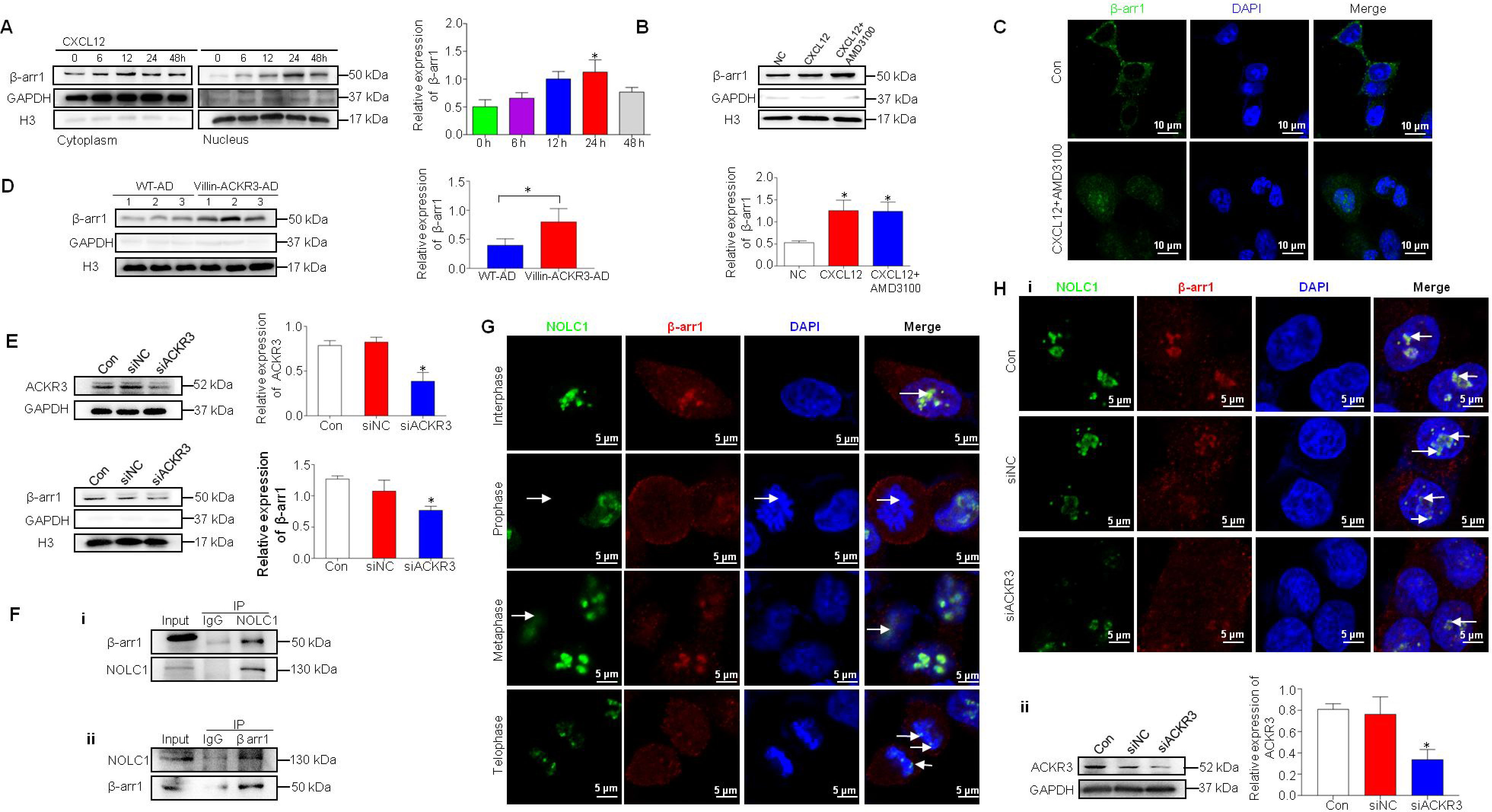
Activation of ACKR3 induced β-arr1 recruitment to the nucleus, interacting with NOLC1. (A) HCT116 cells exposed to CXCL12 (40 ng/ml) for the indicated times, cytoplasmic and nuclear extracts were subjected to Western blotting assay for analysis of β-arr1. The average of nuclear β-arr1 in the graphics corresponds to the quantification of protein bands normalized to H3. *n* = 3. (B) HCT116 cells were exposed to CXCL12 (40 ng/ml) and CXCL12 (40 ng/ml) + AMD3100 (10 μM) for 24 h, nuclear extracts were subjected to Western blotting assay for analysis of β-arr1. *n* = 3. (C) Immunofluorescence staining analyzed the localization of β-arr1 in HCT116 cells exposed to CXCL12 and AMD3100. Scale bar: 10 μm. (D) Western blotting analysis showed the expression of β-arr1 in the nucleus in colorectal cancer tissues of Villin-ACKR3-AD and WT-AD mice. 1, 2, 3 represented the samples from three mice. *n* = 3. (E) Knockdown of ACKR3 reduced the expression levels of β-arr1 and NOLC1. *n* = 3. (F) Co-IP analysis indicated the interaction between β-arr1 and NOLC1. (G) Immunofluorescence analysis showed the variation of NOLC1 and β-arr1 colocalization during cell cycles in HCT116 cells exposed to CXCL12 (40 ng/ml). Scale bar: 5 μm. (H) Knockdown of ACKR3 reduced the colocalization of NOLC1 and β-arr1. Scale bar: 5 μm. Arrows showed the colocalization between NOLC1 and β-arr1. \**P* < 0.05. WT -AD: wild mice were exposed to AOM/DSS. Villin-ACKR3-AD: Villin-ACKR3 mice were exposed to AOM/DSS.

As β-arr1 was translocated into the nucleus, we next investigated the functions of nuclear β-arr1. We had searched any information regarding the interaction of β-arr1 with nuclear proteins issued in the HitPredict database. We observed a specific indication that nuclear β-arr1 might interact with NOLC1. Herein, a strong interaction of nuclear β-arr1 with NOLC1 was identified in HCT116 cells with activated ACKR3 by CXCL12 (Fig 3F, i and ii). Since NOLC1 levels varied during the cell cycles (Pai et al., 1995), we thus identified the interaction of nuclear β-arr1 with NOLC1 more obviously in the interphase and the telophase than in the prophase and the metaphase (Fig 3G). Knockdown of ACKR3 significantly prevented the interaction of nuclear β-arr1 with NOLC1 (Fig 3H).

### The interaction of nuclear β-arr1 with NOLC1 resulted in the promotion of NOLC1 in the nucleolus

NOLC1 is a highly phosphorylated nucleolus protein functions as a regulator of RNA polymerase I (Kim *et al.*, 2003; Meier, 1996). Still little has been known its roles in tumorigenesis. The Cancer Genome Atlas (TCGA) reported high levels of NOLC1 in colorectal cancers (Fig 4A). Our results showed the increased NOLC1 levels in colorectal cancer grown in Villin-ACKR3 mice (Fig 4B) as well as human colonic cancer tissues (Fig 4C). We performed the Immunofluorescence assay to mark the colocalization of NOLC1 with Fibrillarin, the rRNA methyl-transferase, and UBF1, the nucleolar transcription factor 1. NOLC1 was exactly identified in the nucleolus (Fig 4D). Reversely, NOLC1 was not presented in the nucleolus of cancer cells knockdown of β-arr1 (Fig 4E) or ACKR3 (Fig 4F). These results indicated that activation of ACKR3 resulted in the increase of NOLC1 in the nucleolus through the induction of nuclear translocation of β-arr1.

**Figure 4.**
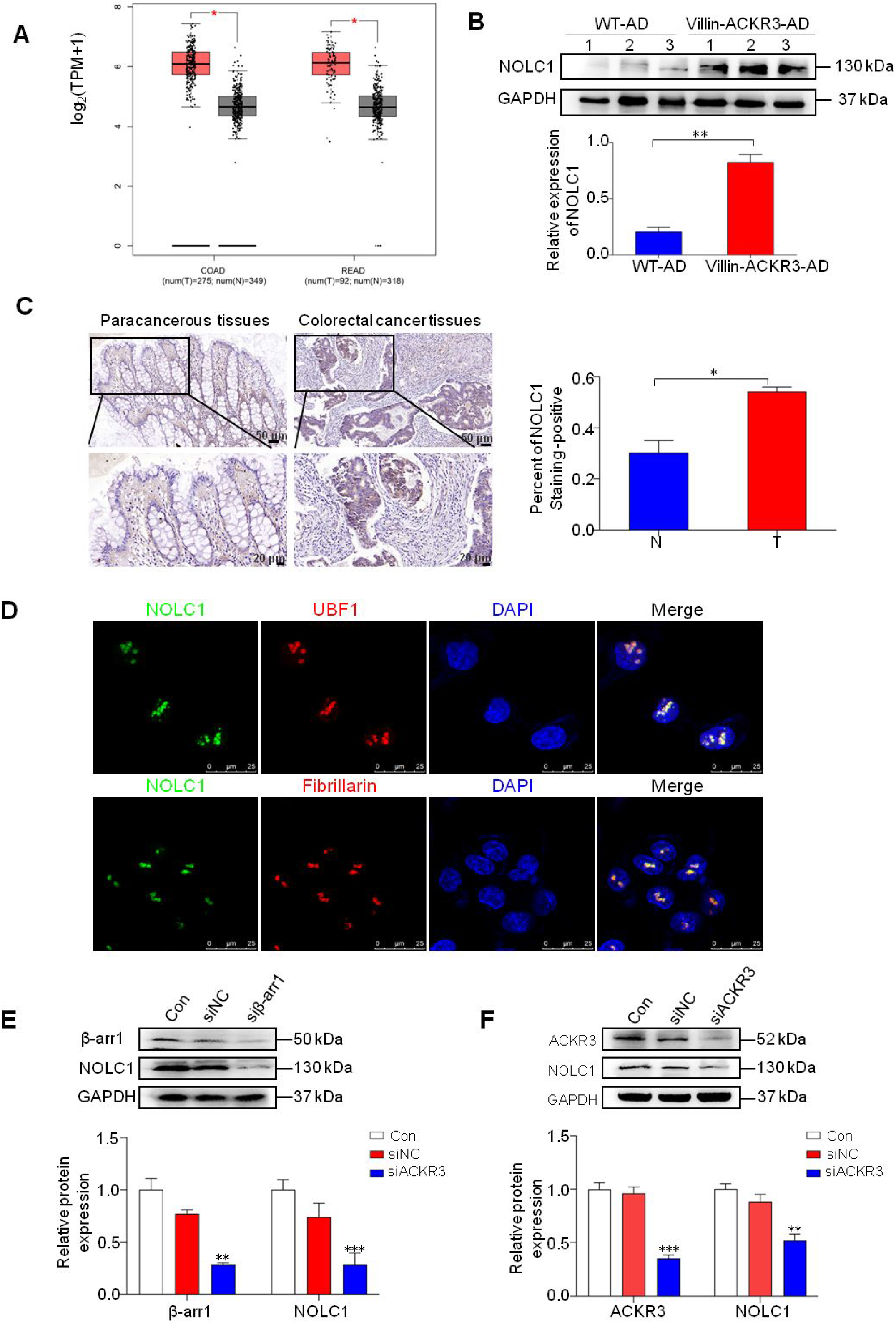
The interaction of nuclear β-arr1 with NOLC1 resulted in the promotion of NOLC1 in the nucleolus. (A) The expression levels of NOLC1 in CRC in TCGA datasets. Differences were seen in NOLC1 levels between colonic tumor (Orange) and corresponding paracancerous tissues (Grey). COAD: Colon adenocarcinoma, READ: Rectum adenocarcinoma. Log2(TPM + 1) was used for log-scale. Calculated means ± SEM were represented by bars and whiskers. (B) Western blotting analysis of NOLC1 levels in Villin-ACKR3-AD and WT-AD mice. 1, 2, 3 represented the samples from three mice. *n* = 3. (C) Expression of NOLC1 in human CRC tissues (marked by T) and paired adjacent non-cancer tissues (marked by N). Upper images 20x, lower images 40x. The boxed region in each top panel is magnified and shown in corresponding bottom panel. (20x) Scale bar: 50 μm, (40x) Scale bar: 20 μm. *n* = 5. (D) NOLC1 was localized to the nucleolus. Co-staining of NOLC1 (Alexa Flour 488), Fibrillarin and UBF1 (Alexa Flour 647) in HCT116 cells. Scale bar: 25 μm. *n* = 3. (E) Knockdown of β-arr1 reduced the expression of NOLC1 in CRC cells. *n* = 3. (F) Western blotting analysis shows the expression levels of NOLC1 in cancer cells with knockdown ACKR3. \**P* < 0.05, \*\**P* < 0.01, \*\*\**P* < 0.001. WT-AD: wild mice exposed to AOM/DSS. Villin-ACKR3-AD: Villin-ACKR3 mice exposed to AOM/DSS.

### The increased nucleolus NOLC1 promoted the synthesis of rRNA of ribosome biogenesis

To investigate the nucleolus NOLC1-promoted in tumorigenesis, we analyzed the level of nuclear AgNOR protein, a marker of cell proliferation, in colorectal cancer tissues of Villin-ACKR3 mice. More deeply and globally staining of AgNOR proteins was measured in colorectal cancer of Villin-ACKR3 than that of WT mice (Fig 5A). To illustrate the role of NOLC1 in the nucleolus, we employed cell model silenced NOLC1 to analyze the levels of synthesis of rRNA of ribosome biogenesis. We designed 3 types of siRNAs to target NOLC1. The levels of NOLC1 were significantly reduced by 51.2%, 49.7%, 65.1%, respectively, in si1-, si2-, and si3-treated HCT116 cells (Fig 5B). Silence of NOLC1 by si3 resulted in a significant reduction of nucleoli (Fig 5C). We then analyzed the levels of precursor rRNA 45S, 36S, and 32S rRNA by RT-qPCR assay. As shown in Fig 5D, the expression levels of pre-rRNA 45S, 36S, and 32S were down-regulated in cancer cells with silenced NOLC1 (siNOLC1 cells) as compared with control WT cells. These results indicated that silencing gene of NOLC1 resulted in the reduction of rRNA synthesis. We further analyzed the nucleolar size of cancer cells with silenced NOLC1 under the transmission electron microscopy. Silencing of NOLC1 gene resulted in the nucleolus smaller than control cells (Fig 5E). Western blotting analysis demonstrated a higher level of nucleolar NOLC1 in colonic cancer tissues in Villin-ACKR3-AD than WT-AD mice. These colonic cancer tissues with elevated NOLC1 exhibited higher level of POLR1A (RPA194), the largest subunit of RNA Pol I, and higher level of UBF1, the transcription initiation factor of rRNA transcription, in Villin-ACKR3-AD than WT mice (Fig 5F). These results suggest that the ACKR3-activated nucleolar NOCL1 is associated with the synthesis of rRNA of ribosome biogenesis.

**Figure 5.**
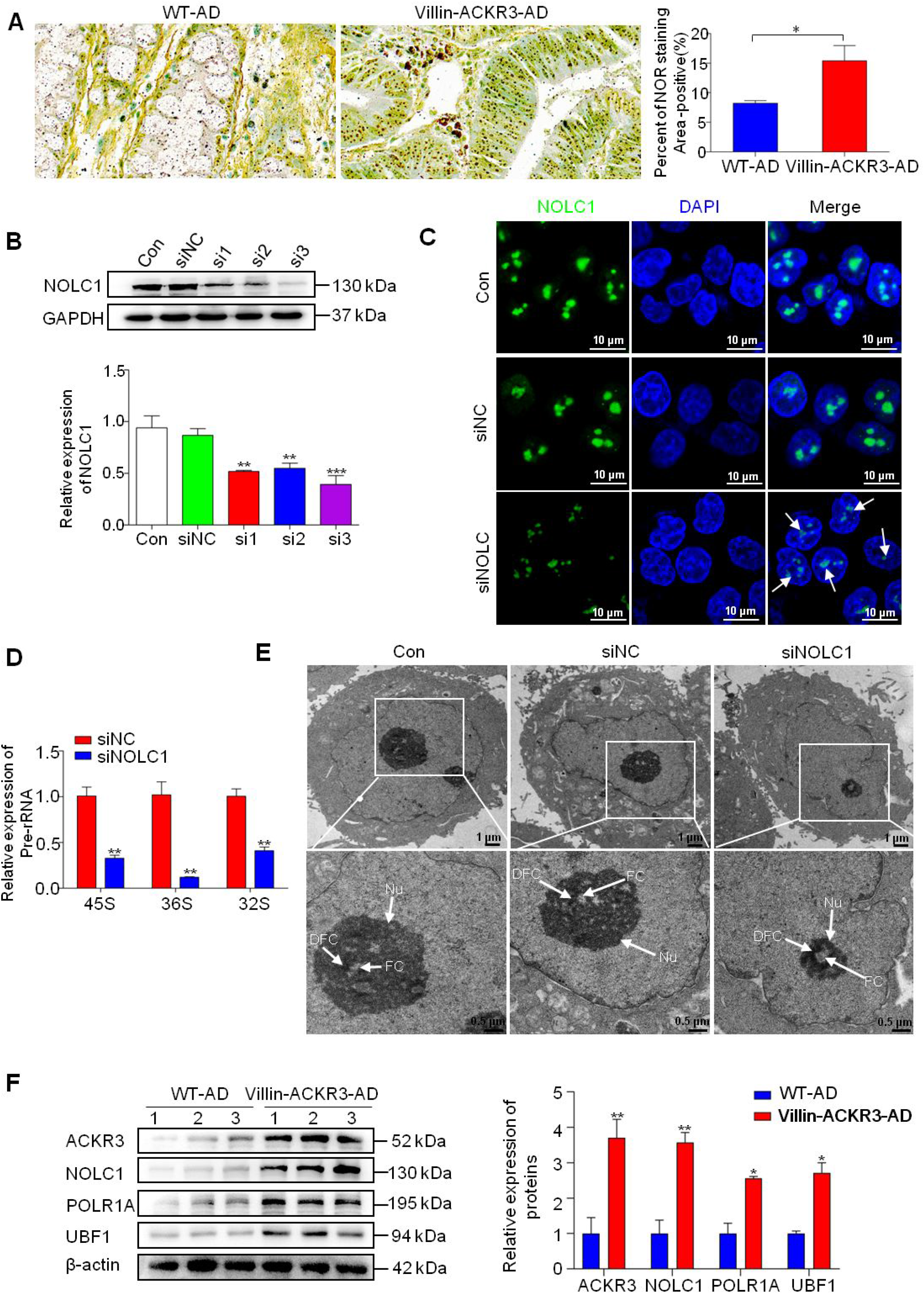
Nuclear β-arr1-activated NOLC1 promoted the synthesis of rRNA of ribosome biogenesis. (A) AgNOR staining of colonic tissues in WT-AD and Villin-ACKR3-AD mice. Scale bar: 10 μm. *n* = 6 (B) Western blotting analysis of NOLC1 levels in HCT116 cells with knockdown ACKR3. *n* = 3 (C) Immunofluorescence staining of NOLC1 in cancer cells. Scale bar: 10 μm. *n* = 3. (D) Real-time PCR analyzed the relative levels of pre-RNA 45S, 36S, and 32S in HCT116 cells. *n* = 3 (E) Transmission electron microscopy (TEM) showed the nucleolus in HCT116 cells of control group (Con), siNC group cells (siNC), silenced NOLC1 group cell (siNOLC1). *n* = 3. (F) Western blotting analysis of ACKR3, NOLC1, POLR1A and UBF1 in colonic tissues of WT-AD and Villin-ACKR3-AD mice. 1, 2, 3 represented the samples from three mice. *n* = 3. DFC, FC and nucleolus were indicated with arrows. \*\**P* < 0.01, \*\*\**P* < 0.001. DFC: Dense Fibrillar Component. FC: Fibrillar Center. Nu: Nucleolus. WT-AD: wild mice exposed to AOM/DSS. Villin-ACKR3-AD: Villin-ACKR3 mice exposed to AOM/DSS.

### The interaction of nucleolar NOLC1 with Fibrillarin led to the increase of Fibrillarin and resulted in the promotion of rRNA transcription

Fibrillarin plays an important role in tumorigenesis through inducing rRNA transcription (Shubina et al., 2018). As we identified the function of nucleolar NOLC1 in promoting synthesis of rRNA of ribosome biogenesis, we wanted to know whether Fibrillarin was involved in the NOCL1-activated synthesis of rRNA. Analysis of correlation data issued by GEPIA database indicated a correlation of nucleolar NOLC1 to Fibrillarin in CRC tissues (correlation coefficient R = 0.4, Fig 6A). Further immunofluorescence analysis indicated a strong colocalization of NOLC1 with Fibrillarin in nucleolus of HCT116 cells (Fig 6B). Western blotting analysis indicated a significant higher Fibrillarin in Villin-ACKR3 mice than WT mice (Fig 6C). Conversely, knockdown of nucleolar NOLC1 significantly reduced Fibrillarin levels (Fig 6D). The interaction of nucleolar NOLC1 with Fibrillarin was clearly seen in colorectal cancer cells of Villin-ACKR3 mice but not obviously in colorectal cancer cells of WT mice (Fig 6E). These results indicated that the interaction of NOLC1 with Fibrillarin induced the upregulation of Fibrillarin and further promoted rRNA transcription in colorectal cancer cells of Villin-ACKR3 mice. We conclude that activation of ACKR3 promotes colorectal tumorigenesis through the increase of ribosome biogenesis by the interaction of nuclear NOLC1 with Fibrillarin (Fig 6F).

**Figure 6.**
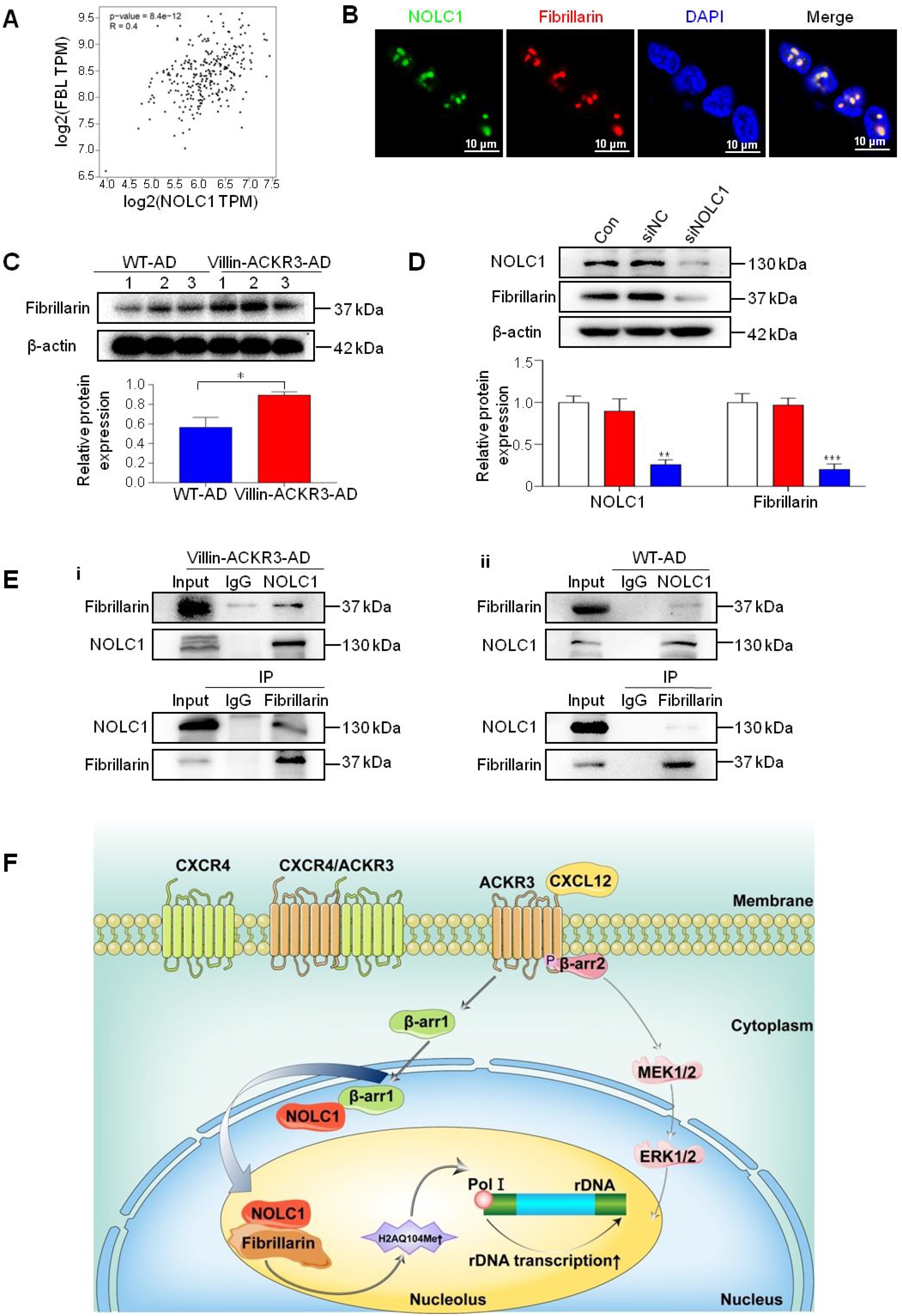
The interaction of NOLC1 with Fibrillarin led to the increase of Fibrillarin and resulted in the promotion of rRNA transcription. (A) The correlation of NOLC1 to Fibrillarin in human colorectal cancer tissues in GEPIA database. (B) Immunofluorescence staining of NOLC1 interacted with Fibrillarin in the nucleolus. Scale bar: 10 μm. *n* = 3. (C) The expression of Fibrillarin in colorectal cancer tissues of Villin-ACKR3-AD and WT-AD mice. 1, 2, 3 represented the samples from three mice. *n* = 3. (D) Knockdown of NOLC1 reduced Fibrillarin level in HCT116 cells. *n* = 3. (E) Co-IP analysis indicated an interaction between NOLC1 and Fibrillarin. (F) Proposed mechanism of activated ACKR3 in colorectal tumorigenesis. Activation of ACKR3 induces nuclear translocation of β-arr1, leading to the interaction of NOLC1, resulting in the Fibrillarin-induced rRNA transcription of ribosome biogenesis. \**P* < 0.05, \*\**P* < 0.01, \*\*\**P* < 0.001. WT-AD: wild mice exposed to AOM/DSS. Villin-ACKR3-AD: Villin-ACKR3 mice exposed to AOM/DSS.

## Discussion

It is known that, as an atypical GPCR, targeting ACKR3 does not lead to typical G-protein-coupled receptor-mediated calcium mobilization and chemotaxis, but rather the recruitment of β-arrestins and the internalization of GPCR (Nguyen et al., 2020). ACKR3 has long been considered as a scavenger receptor and decoy receptor (Luker *et al.*, 2012; Meyrath *et al.*, 2020). ACKR3 is upregulated in the inflammatory cells and the malignant cells. Even though ACKR3 was found to function in the ACKR3/CXCR4 heterodimer in tumorigenesis (Koch and Engele, 2020), much less is known about the mechanism of ACKR3. Since Villin-ACKR3 mice developed more exacerbated colorectal cancer than Villin-CXCR4 mice, thus we hypothesize that ACKR3 play a crucial role independent of CXCR4. In the present study, our clinical data showed that high ACKR3 was associated with increased severity of clinical stages. ACKR3 has been received attention in diagnostics, targeting drug design, and management of patients.

To unveil the mechanism of ACKR3 in colorectal tumorigenesis, we firstly answered the question that how the activated ACKR3 signaling be translocated into the cytoplasm and then to the nucleus. We have understood that GPCRs could interact with β-arrestins (β-arr1 and β-arr2) to function as scaffolds for a multiple kinases that connect GPCRs to the effector pathways. These complex signaling network and interactions are facilitated by a conformational change in the β-arrs that is thought to occur upon binding to a phosphorylated activated GPCR (Charest et al., 2005). Previous reports indicated that activation of ACKR3 could recruit the β-arr2 into the nucleus through the AKT and ERK1/2 signaling pathways (Li et al., 2014; Lin et al., 2014; Min et al., 2020). Since β-arr2 has a nuclear export signal (NES) in its C terminus, however, this specific NES excludes it from sustained presence in the nucleus (Scott *et al.*, 2002). In the present study, we revealed that activation of ACKR3 induced nuclear translocation of β-arr1 but not β-arr2 into the nucleus. Importantly, the ACKR3-induced β-arr1 interacted with NOLC1, a most highly nuclear phosphoprotein, to form the β-arr1-NOLC1 complex.

How NOLC1 was identified as the downstream effector of nuclear β-arr1? We firstly searched for the clues about the interaction related to the nuclear β-arr1 in the Hitpredict database. An interesting link of β-arr1 to NOLC1 was found in the database. Previously, NOLC1 was already identified as a nuclear localization signal-binding protein and functions as a chaperone for shuttling between the nucleolus and the cytoplasm (Meier and Blobel, 1992). However, the underlying mechanisms and biological functions remain largely unknown, and its roles in tumorigenesis are contradictious among the reports (Duan et al., 2013; Huang et al., 2018; Hwang et al., 2009; Yuan et al., 2017b). Since NOLC1 was upregulated in the interphase and the telophase, it is thus suggested that the interaction of nuclear β-arr1 with NOLC1 might play the function of over-proliferation in cancer cells (Pai *et al.*, 1995). However, there were some confused reports that nuclear NOLC1 increased the percentage of cells in their G0/G1 phase (Yuan et al., 2017a). Our further research revealed that the ACKR3-induced interaction of β-arr1 with NOLC1 exactly occurred in the interphase and the telophase but not in the prophase and the metaphase. These results indicated that the interaction of β-arr1 with NOLC1 resulted in the upregulation of NOLC1 in the nucleolus.

We next identified downstream effector of nuclear NOLC1. High ribosome biogenesis has long been considered to associate with key process of over-proliferation and tumorigenesis in many types of cells (Penzo et al., 2019). Ribosome biogenesis is a universal, complex and well-orchestrated cellular process, such as production of ribosomal RNA, synthesis of ribosomal proteins and ribosome assembly (Pelletier et al., 2018). Regulation of ribosome biogenesis, particularly the production of rRNA, is a critical aspect of cell growth control (Mayer and Grummt, 2006). Nucleolus, a nuclear subcompartment where ribosomal RNA is synthesized and assembled into the ribosomal subunits, is the main site of ribosome biogenesis (Grummt, 2013). It is a dynamic organelle subject to inputs from growth signaling pathways, nutrients, and stress, whose size correlates with the rRNA synthesis (Boulon et al., 2010). NOLC1 interacts with RNA polymerase I, leading to the regulation of rRNA transcription, perhaps through linking RNA Pol I transcription with pre-rRNA processing. NOLC1 also plays an essential role in rDNA transcription and further inducing rRNA in ribosome biogenesis (Chen et al., 1999; Meier, 2005). Thus, ribosome biogenesis is identified as the downstream effector of nuclear NOLC1. In the present study, we found that colorectal cancer grown in Villin-ACKR3 mice demonstrated increased ribosome biogenesis, and knockdown of NOLC1 resulted in a decrease of pre-rRNA and smaller nucleolar size in cancer cells. It is thus that increased NOLC1 functions a key process in synthesis of rRNA. Increased NOLC1 was found to further interact with Fibrillarin in the process of tumorigenesis. Colorectal cancer developed in Villin-ACKR3 mice demonstrated a strong interaction of NOLC1 to Fibrillarin in the nucleolus. Conversely, knockdown of NOLC1 downregulated the expression levels of Fibrillarin in the nucleolus. Fibrillarin is a highly conserved nucleolus protein. Fibrillarin plays a crucial role in the regulation of ribosome biogenesis through promoting the methylation of ribosomal RNAs and rDNA histones (El Hassouni et al., 2019). Accordingly, Fibrillarin might be a downstream signaling of NOLC1. However, the mechanism of NOLC1-activated Fibrillarin in the ribosome biogenesis has not been investigated. A report indicated that enhanced Fibrillarin might increase rRNA transcription through activating RNA Pol I methylation in H2AQ104 (Tessarz et al., 2014). In colorectal cancer of Villin-ACKR3 mice, ACKR3-activated NOLC1 promoted nuclear Fibrillarin expression, accordingly, enhanced Fibrillarin functions the upregulation of ribosome biogenesis through promoting methylation of ribosomal RNAs and rDNA histones.

In conclusion, ACKR3 promotes colorectal tumorigenesis through the perturbation of rRNA biogenesis by nuclear β-arr1-induced interaction of NOLC1 with Fibrillarin. We envision that future approaches to treat colorectal cancer should use ACKR3 inhibitors for preventing the ACKR3-activated NOLC1 and Fibrillarin in the nucleolus.

## Materials and Methods

### TCGA dataset

We referenced the microarray databases of Oncomine (http://www.oncomine.org) (Rhodes et al., 2004) and GEPIA (http://gepia.cancer-pku.cn) (Tang et al., 2017) for analyzing the expression of ACKR3 and NOLC1 in cancer tissues and normal tissues. To analyze the dataset, thresholds were set as follows: p-value: 0.01; fold change: 2; gene rank: 20%; analysis type: cancer *vs* normal; data type: mRNA.

### Human colonic cancer specimens

Human CRC specimens and their paired adjacent non-neoplastic tissues were collected from resected specimens between February and May of 2018 in Binzhou Medical College Hospital (Binzhou, China) (n = 20). The use of human clinical samples was approved by the Ethics Committee of Binzhou Medical College.

### Human colon cell lines, cell culture and treatment

Human cancer cell lines SW480, SW620, HCT116, HT29 and human normal colonic cell line NCM460 were purchased from China Cell Bank authorized by American Type Culture Collection (ATCC). Cells were cultured in medium of RPMI1640 or DMEM (Gibco) containing 10% FBS (Gibco) at 37°C in a humid atmosphere (5% CO_2_). Cells were exposed to 40 ng/ml SDF-1α (CXCL12) (PeproTech) and AMD3100 10 μM (Selleck) for analysis of ACKR3 expression and its downstream signaling pathways.

### Villin-ACKR3 mice and establishment of colorectal cancer

Villin-ACKR3-IREF mice specifically expressed high ACKR3 in intestinal epithelial cells (IECs) were generated by Cyagen Biosciences Inc (Guangzhou, China). Villin-ACKR3-IREF mice and their WT littermates were maintained under controlled room temperature and allowed unrestricted access to the standard mouse feed (Meidenbauer et al., 2014). Animal experiments were approved by Animal Welfare Committee of Capital Medical University (permit no. AEEI-2016043). Mice model of colorectal cancer was established as previously described (Song *et al.*, 2019).

### Histopathology and Immunohistochemical staining assay

Routine hematoxylin-eosin staining (H&E staining) of 5 μm thick sections were used for histological analysis of cancer tissues. Immunohistochemical staining assay (IHC) was performed using antibodies against ACKR3 (Abcam), PCNA (Proteintech Group), NOLC1 (Proteintech Group). IHC scoring was obtained based on the staining intensity and percentage of stained cancer cells as described elsewhere.

### RNA interference assay

Small interfering RNA (siRNA) was designed by GenePharma (Suzhou, China). Human siACKR3, siβ-arr1, siNOLC1 were co-transfected into cells using Lipofectamine 2000 (Invitrogen) (Sequences of siRNAs were detailed in Supplementary Table S1). Cells were plated in the 6-well plates for 24 h before interfering. The diluted Lipofectamine 2000 was lightly mixed with an equal volume siRNA suspension. Each sample was incubated at room temperature for 20 min. The siRNA-lipid complex was added and incubated at 37°C for 6 h. Finally, new medium was replaced and cells were incubated for 48 h before treatment.

### RNA extraction and quantitative PCR (real-time PCR)

Total RNA was extracted using TRIzol reagent according to manufacturer’s protocols (Invitrogen). Reverse transcription of total RNA, the first-strand cDNA was synthesized with ReverTra Ace^®^ qPCR RT Kit (TOYOBO, Japan). Quantitative PCR was performed in triplicate using SYBR^®^ Green Realtime PCR Master Mix (TOYOBO, Japan) on an ABI 7500 Real Time PCR System. β-actin (ATCB) was used as an endogenous control, and fold changes were calculated by means of relative quantification (2^−ΔΔ^CT). The primers used for quantitative PCR were listed in Supplementary Table S2.

### Total protein and nuclear protein extraction

Total protein was extracted using a strong version of RIPA lysate (Beyotime Biotechnology, China), containing 1 mM PMSF. Nuclear protein was isolated using NE-PER nuclear and cytoplasmic extraction reagent (Beyotime Biotechnology, China) according to manufacturer’s protocols. Protein concentrations were quantitatively determined using BCA protein assay kit (Beyotime Biotechnology, China).

### Western blotting assay

Western blotting assay was performed as described previously (Zhang et al., 2020). Primary antibodies were listed in Supplementary Table S3. Appropriate horseradish peroxidase-conjugated secondary antibodies were purchased from ZSGB-BIO, China. Images were obtained by FluorChem FC3 image analyzer (Molecular Devices). Band intensity was analyzed with ImageJ analysis program.

### Co-immunoprecipitation (CO-IP)

Samples of cultured cells and cancer tissues were obtained by NP-40 lysis buffer (Beyotime Biotechnology, China), containing 1 mM PMSF. Complexes of proteins, protein A/G beads and specific antibodies used Beaver Beads™ Protein A/G Immunoprecipitation kit (Beaver Biosciences Inc, China) according to manufacturer’s protocols. Immunoprecipitates were resolved by 10% SDS-PAGE membranes (Millipore). Membranes were incubated in blocking buffer [1% (w/v) BSA, 5% (w/v) non-fat dry milk, and 0.1% (v/v) Tween-20 in TBS (pH 7.0)] overnight at room temperature. Membranes were subsequently probed with the corresponding antibodies in blocking buffer for 12-16 h at 4°C. These antibodies included NOLC1 polyclonal antibody (Proteintech Group), β-arrestin1 monoclonal antibody (Invitrogen) and Fibrillarin antibody (Santa Cruz). Membranes were incubated with HRP-conjugated anti-rabbit IgG or HRP-conjugated anti-mouse IgG for 1 h at room temperature. Antibody-reactive proteins were detected by western chemiluminescent HRP substrate (ECL) (Millipore). Images were acquired by FluorChem FC3 image analyzer (Molecular Devices).

### Immunofluorescence assay

Cells (3-5×10^4^) cultured onto glass were fixed in 100% cold methanol for 10 min, washed in PBS, permeabilized with 0.5% Triton (Sigma-Aldrich) for 30 min, blocked for nonspecific antibody reactions by incubating in solution containing 5% bovine serum albumin for 30 min, incubated with primary antibodies of β-arrestin1 anti-rabbit IgG (Abcam), Fibrillarin (Abcam), NOLC1 (Proteintech Group), β-arrestin1 anti-mouse IgG (Invitrogen), UBF1 (Santa Cruz) at 4°C overnight, followed by staining with secondary antibodies of Alexa Fluor™ 488 donkey anti-rabbit IgG (Invitrogen) or Alexa Fluor^®^647 goat anti-mouse IgG (Cell Signaling Technology) for 1 h at room temperature. Cell nuclei were counterstained with 4’6-diamidino-2-phenylindole (DAPI, Beyotime Biotechnology, China). Images were acquired under confocal laser imaging system (NIS-Elements Confocal, Japan).

### AgNOR staining

The method of AgNOR staining was performed as described previously (Trerè, 2000). Paraffin sections deparaffinized were washed for 10 min and then stained in AgNOR Stain solution (Solarbio, China) at room temperature for 40-60 min. After rinsing in distilled water for 1-2 min, slides were counterstained for 1-3 min in methyl green staining solution, washed and finally air dried. AgNOR staining was analyzed by area quantification module of HALO v3.0.311.407 (Indica Labs) software.

### Transmission electron microscopy (TEM)

Sample preparation for TEM, cancer cells were aspirated into the centrifuge tube and centrifuged at 800 rpm for 5 min. The cell pellets were fixed by adding 2.5% glutaraldehyde (Servicebio, China) for 2 h at 4°C. Cell pellets were rinsed three times and then fixed with 1% osmium tetroxide for 1 h. After dehydration through a graded ethanol series, samples were embedded in propylene oxide and resin (1:1). Sections were examined with a transmission electron microscope (JEM-2100, Japan).

### Statistical analysis

Data were described as mean ± SEM. Statistical analysis was done with GraphPad Prism 8. Student’s t test was used to compare differences between two groups. Statistical differences among multiple groups were analyzed by one-way analysis of variance (ANOVA) followed by post hoc test with Dunnett (multiple comparisons to the same control). *P* values < 0.05 were considered significant.

## Acknowledgement

This work was supported by National Natural Science Foundation of China (81973350/81872884/81772637) and Beijing Natural Science Foundation (7212149).

## Author contributions

S-X Cui conceived and designed the study. J Yang, Y-N Li, T Pan, R-R Miao and Y-Y Zhang performed the experiments. S-H Wu provided human colonic cancer specimens. J Yang wrote the manuscript and performed statistical analysis. X-J Qu provided intellectual inputs and edited the manuscript.

## References

Bachelerie, F., Graham, G.J., Locati, M., Mantovani, A., Murphy, P.M., Nibbs, R et al (2014) New nomenclature for atypical chemokine receptors. Nat Immunol 15: 207–208

Balabanian, K., Lagane, B., Infantino, S., Chow, K.Y., Harriague, J., Moepps, B et al (2005) The chemokine SDF-1/CXCL12 binds to and signals through the orphan receptor RDC1 in T lymphocytes. J Biol Chem 280: 35760–35766

Becker, J.H., Gao, Y., Soucheray, M., Pulido, I., Kikuchi, E., Rodriguez, M.L et al (2019) CXCR7 reactivates ERK signaling to promote resistance to EGFR kinase inhibitors in NSCLC. Cancer Res 79: 4439–4452

Boulon, S., Westman, B.J., Hutten, S., Boisvert, F.M., Lamond, A.I (2010) The nucleolus under stress. Mol Cell 40: 216–227

Burns, J.M., Summers, B.C., Wang, Y., Melikian, A., Berahovich, R., Miao, Z et al. (2006) A novel chemokine receptor for SDF-1 and I-TAC involved in cell survival, cell adhesion, and tumor development. J Exp Med 203: 2201–2213

Cancellieri, C., Vacchini, A., Locati, M., Bonecchi, R., Borroni, E.M (2013) Atypical chemokine receptors: from silence to sound. Biochem Soc Trans 41: 231–236

Charest, P.G., Terrillon, S., and Bouvier, M (2005) Monitoring agonist-promoted conformational changes of beta-arrestin in living cells by intramolecular BRET. EMBO Rep 6: 334–340

Chen, H.K., Pai, C.Y., Huang, J.Y., Yeh, N.H (1999) Human Nopp140, which interacts with RNA polymerase I: implications for rRNA gene transcription and nucleolar structural organization. Mol Cell Biol 19: 8536–8546

Duan, X., Zhang, J., Liu, S., Zhang, M., Wang, Q., Cheng, J (2013) Methylation of nucleolar and coiled-body phosphoprotein 1 is associated with the mechanism of tumorigenesis in hepatocellular carcinoma. Oncol Rep 30: 2220–2228

El Hassouni, B., Sarkisjan, D., Vos, J.C., Giovannetti, E., Peters, G.J (2019) Targeting the ribosome biogenesis key molecule Fibrillarin to avoid chemoresistance. Curr Med Chem 26: 6020–6032

Grummt, I (2013) The nucleolus-guardian of cellular homeostasis and genome integrity. Chromosoma 122: 487–497

Huang, H., Li, T., Chen, M., Liu, F., Wu, H., Wang, J (2018) Identification and validation of NOLC1 as a potential target for enhancing sensitivity in multidrug resistant non-small cell lung cancer cells. Cell Mol Biol Lett 23: 54

Hwang, Y.C., Lu, T.Y., Huang, D.Y., Kuo, Y.S., Kao, C.F., Yeh, N.H (2009) NOLC1, an enhancer of nasopharyngeal carcinoma progression, is essential for TP53 to regulate MDM2 expression. Am J Pathol 175: 342–354

Kim, Y.K., Jin, Y., Vukoti, K.M., Park, J.K., Kim, E.E., Lee, K.J (2003) Purification and characterization of human nucleolar phosphoprotein 140 expressed in Escherichia coli. Protein Expr Purif 31: 260–264

Koch, C., Engele, J (2020) Functions of the CXCL12 Receptor ACKR3/CXCR7-What has been perceived and what has been overlooked. Mol Pharmacol 98: 577–585

Li, S., Fong, K.W., Gritsina, G., Zhang, A., Zhao, J.C., Kim, J., Sharp, A et al (2019) Activation of MAPK signaling by CXCR7 leads to enzalutamide resistance in prostate cancer. Cancer Res 79: 2580–2592

Li, X.X., Zheng, H.T., Huang, L.Y., Shi, D.B., Peng, J.J., Liang, L et al (2014) Silencing of CXCR7 gene represses growth and invasion and induces apoptosis in colorectal cancer through ERK and beta-arrestin pathways. Int J Oncol 45: 1649–1657

Lin, L., Han, M.M., Wang, F., Xu, L.L., Yu, H.X., Yang, P.Y (2014) CXCR7 stimulates MAPK signaling to regulate hepatocellular carcinoma progression. Cell Death Dis 5: e1488

Luker, K.E., Lewin, S.A., Mihalko, L.A., Schmidt, B.T., Winkler, J.S., Coggins, N.L et al (2012) Scavenging of CXCL12 by CXCR7 promotes tumor growth and metastasis of CXCR4-positive breast cancer cells. Oncogene 31: 4750–4758

Mayer, C., and Grummt, I (2006) Ribosome biogenesis and cell growth: mTOR coordinates transcription by all three classes of nuclear RNA polymerases. Oncogene 25: 6384–6391

Meidenbauer, J.J., Ta, N., Seyfried, T.N (2014) Influence of a ketogenic diet, fish-oil, and calorie restriction on plasma metabolites and lipids in C57BL/6J mice. Nutr Metab (Lond) 11: 23

Meier, U.T (1996) Comparison of the rat nucleolar protein Nopp140 with its Yeast Homolog SRP40. J Biol Chem 271: 19376–19384

Meier, U.T (2005) The many facets of H/ACA ribonucleoproteins. Chromosoma 114: 1–14

Meier, U.T., Blobel, G (1992) Nopp140 shuttles on tracks between nucleolus and cytoplasm. Cell 70: 127–138

Meyrath, M., Szpakowska, M., Zeiner, J., Massotte, L., Merz, M.P., Benkel, T et al. (2020) The atypical chemokine receptor ACKR3/CXCR7 is a broad-spectrum scavenger for opioid peptides. Nat Commun 11: 3033

Miao, Z., Luker, K.E., Summers, B.C., Berahovich, R., Bhojani, M.S., Rehemtulla, A et al (2007) CXCR7 (RDC1) promotes breast and lung tumor growth in vivo and is expressed on tumor-associated vasculature. Proc Natl Acad Sci USA 104: 15735–15740

Min, K., Yoon, H.J., Park, J.Y., Baidya, M., Dwivedi-Agnihotri, H., Maharana, J et al (2020) Crystal structure of beta-arrestin 2 in complex with CXCR7 phosphopeptide. Structure 28: 1014–1023

Nguyen, H.T., Reyes-Alcaraz, A., Yong, H.J., Nguyen, L.P., Park, H.K., Inoue, A et al (2020) CXCR7: a beta-arrestin-biased receptor that potentiates cell migration and recruits beta-arrestin2 exclusively through Gbetagamma subunits and GRK2. Cell Biosci 10: 134

Pai, C.Y., Chen, H.K., Sheu, H.L., Yeh, N.H (1995) Cell-cycle-dependent alterations of a highly phosphorylated nucleolar protein p130 are associated with nucleologenesis. J Cell Sci 108 ( Pt 5): 1911–1920

Pelletier, J., Thomas, G., Volarević, S (2018) Ribosome biogenesis in cancer: new players and therapeutic avenues. Nat Rev Cancer 18: 51–63

Penzo, M., Montanaro, L., Trere, D., Derenzini, M (2019) The ribosome biogenesis-cancer connection. Cells 8: 5

Rajagopal, S., Kim, J., Ahn, S., Craig, S., Lam, C.M., Gerard, N.P et al (2010) Beta-arrestin-but not G protein-mediated signaling by the “decoy” receptor CXCR7. Proc Natl Acad Sci USA 107: 628–632

Rhodes, D.R., Yu, J., Shanker, K., Deshpande, N., Varambally, R., Ghosh, D et al (2004) ONCOMINE: a cancer microarray database and integrated data-mining platform. Neoplasia 6: 1–6

Salanga, C.L., O’Hayre, M., Handel, T (2009) Modulation of chemokine receptor activity through dimerization and crosstalk. Cell Mol Life Sci 66: 1370–1386

Sanchez-Martin, L., Sanchez-Mateos, P., Cabanas, C (2013) CXCR7 impact on CXCL12 biology and disease. Trends Mol Med 19: 12–22

Scott, M.G., Le Rouzic, E., Perianin, A., Pierotti, V., Enslen, H., Benichou, S et al (2002) Differential nucleocytoplasmic shuttling of beta-arrestins. Characterization of a leucine-rich nuclear export signal in beta-arrestin2. J Biol Chem 277: 37693–37701

Shubina, M.Y., Musinova, Y.R., and Sheval, E.V (2018) Proliferation, cancer, and aging-novel functions of the nucleolar methyltransferase fibrillarin? Cell Biol Int 42: 1463–1466

Song, Z.Y., Wang, F., Cui, S.X., Gao, Z.H., Qu, X.J (2019) CXCR7/CXCR4 heterodimer-induced histone demethylation: a new mechanism of colorectal tumorigenesis. Oncogene 38: 1560–1575

Sun, X., Cheng, G., Hao, M., Zheng, J., Zhou, X., Zhang, J et al (2010) CXCL12/CXCR4/CXCR7 chemokine axis and cancer progression. Cancer Metastasis Rev 29: 709–722

Tang, Z., Li, C., Kang, B., Gao, G., Li, C., Zhang, Z (2017) GEPIA: a web server for cancer and normal gene expression profiling and interactive analyses. Nucleic Acids Res 45: W98–W102

Tessarz, P., Santos-Rosa, H., Robson, S.C., Sylvestersen, K.B., Nelson, C.J., Nielsen, M.L et al (2014) Glutamine methylation in histone H2A is an RNA-polymerase-I-dedicated modification. Nature 505: 564–568

Trerè, D (2000) AgNOR staining and quantification. Micron 31: 127–131

Wang, J., Shiozawa, Y., Wang, J., Wang, Y., Jung, Y., Pienta, K.J et al (2008) The role of CXCR7/RDC1 as a chemokine receptor for CXCL12/SDF-1 in prostate cancer. J Biol Chem 283: 4283–4294

Yuan, F., Li, G., and Tong, T (2017a) Nucleolar and coiled-body phosphoprotein 1 (NOLC1) regulates the nucleolar retention of TRF2. Cell Death Discov 3: 17043

Yuan, F., Zhang, Y., Ma, L., Cheng, Q., Li, G., Tong, T (2017b) Enhanced NOLC1 promotes cell senescence and represses hepatocellular carcinoma cell proliferation by disturbing the organization of nucleolus. Aging Cell 16: 726–737

Zhang, Y.H., Luo, D.D., Wan, S.B., Qu, X.J (2020) S1PR2 inhibitors potently reverse 5-FU resistance by downregulating DPD expression in colorectal cancer. Pharmacol Res 155: 104717

